# Nanogram of Arg 1 Delivered by extracellular vesicles restore ARG1 activity in a mouse model of ARG1-D and improve lifespan

**DOI:** 10.1101/2025.09.23.677947

**Authors:** Li-En Hsieh, Mafalda Cacciottolo, Michael J. LeClaire, William Morrison, Bailey Murphy, Christy Lau, Kristi Elliott, Linda Marban, Minghao Sun

## Abstract

Arginase 1 (ARG1) deficiency (ARG1-D) is a rare genetic disorder due to loss of ARG1, the final enzyme in the urea cycle. ARG1-D hepatocytes are impaired in converting arginine into urea, resulting in elevated peripheral arginine and ammonia, which leads to progressive neurological symptoms. Current therapeutic strategies mainly focus on managing plasma arginine and ammonia level, but long-term outcomes remain poor. While no approved treatment specific for ARG1-D is available in the United States, a recombinant protein based enzyme replacement therapy is available in Europe. Recently, extracellular vesicles (EVs) are emerging as a powerful therapeutic vehicle. By using Capricor’s StealthX™ platform, EVs were engineered to express human ARG1 on their surface or encapsulated within. Regardless of its localization on the EV membrane, nanograms of ARG1 carried by EVs were biologically active and able to convert arginine into urea as potent as micrograms of human recombinant ARG1 (rHuArg1). Furthermore, ARG1- encapsulating EVs (STX-Arg1-in) were able to deliver ARG1 intracellularly but not EVs carrying ARG1 on their surface or rHuArg1. STX-Arg1-in EVs were further evaluated in a series of in vivo studies and results showed that STX-Arg1-in EVs were non-toxic and able to restore the arginase activities in the liver of the Arg1^-/-^ mice, which leads to a lowered plasma arginine concentration similar to wildtype mice. Most importantly, Arg1-in expanded the life span of the lethal neonatal Arg1 deficiency mouse model. Taken together, our data suggested StealthX™-engineered STX-Arg1-in EVs have a better safety profile due to extreme low dosage and had great potential as a novel enzyme replacement strategy for patients suffering from ARG1-D.

**Significance statement:** Intracellular delivery of recombinant protein and improved life span are endpoints of a successful enzyme replacement therapy for the treatment of ARG1-D. Using StealthX platform, a fully functional ARG1 enzyme was engineered to be carried inside of the extracellular vesicles, which allowed intracellular delivery of ARG1 protein in vitro and in vivo, with improvement of life span in a lethal neonatal mouse model of Arg1 deficiency. More importantly, no toxicity was observed, and efficacy was achieved with low dose, setting the base for an improved therapeutic approach.

## Introduction

Arginase 1 (ARG1) deficiency (ARG1-D) is a rare autosomal recessive genetic disorder that occurs in approximately 2.8 cases per million births with the population prevalence at approximately 1.4 cases per million people worldwide (1, 2). ARG1-D is one of the diseases among the urea cycle disorders (UCDs) and is caused by the mutations or deletions of the *ARG1* gene located on chromosome 6 (6q23) which leads to a lack of ARG1 protein expression in the patients (3). ARG1 is the last enzyme in the urea cycle to catalyze the conversion of arginine into ornithine and urea (3). Urea cycle mainly takes place in hepatocytes in the liver and is the major pathway to detoxify ammonia in mammals (3). Patients with ARG1-D usually display symptoms in their late infancy to pre-school age (4). With the disease progression, patients would show spasticity, mainly in their lower limbs, intellectual disability, motor deficits, seizures, developmental delays/growth deficiency, and other neurological symptoms, which impact the quality of life of the patients and their parents/caregivers (1, 3, 5). The standard therapeutic interventions mainly focused on controlling plasma arginine, through dietary protein restrictions, supplementing arginine-free essential amino acids, and administration of nitrogen scavengers to control the plasma ammonia level (1, 3, 5, 6). Over the past two decades, different approaches for ARG1-D-specific therapies have been evaluated, such as recombinant human ARG1 protein (7), adeno-associated virus (AAV)-based therapies (8–11), and lipid-nanoparticles-based mRNA therapy (12). Despite the promising data in the pre-clinical stage, none of these strategies have produced a valuable therapy but the pegylated human ARG1, which was recently approved in the European Union and United Kingdom (https://www.ema.europa.eu/). Although pegylated human ARG1 treatment resulted in improved plasma arginine level in the pre-clinical studies (7) and clinical trials (13, 14), this approach showed no improvement of lifespans in ARG1-D mouse models (7), presumably due to lack of liver-specific delivery.

For the past decade, extracellular vesicles (EVs) have drawn significant attention for their potential applications in both diagnostics and therapeutics (15, 16). EVs are nano-sized vesicles naturally secreted by almost all cell types (17, 18), of ∼50 – 150 nm in size, formed from the inward budding of the endosome membranes, functioning as “messenger” between cells, involved in different physiological and pathological conditions (19). EVs showed low immunogenicity with good biocompatibility and good safety profile, which are the key features for ideal nanomedicine (15, 19–22). Numerous studies have demonstrated the capability of loading different therapeutic cargos, including protein, small RNA, and small molecules, and many of them are being evaluated in the early to late stages of clinical trials (19, 20, 22, 23). However, so far, there is no approved EV-based therapy according to the US Food and Drug Administrations (https://www.fda.gov/vaccines-blood-biologics/consumers-biologics/consumer-alert-regenerative-medicine-products-including-stem-cells-and-EVs).

The goal of the present study is to develop an enzyme replacement therapy for ARG1-D using our StealthX™ exosome-based platform. We successfully engineered 293F cells to express a functional ARG1 enzyme onto the EVs (STX-Arg1) membrane, which resulted in ARG1 delivery to the cells, while retaining its enzymatic activity. Our data suggested that the ARG1 EVs were safe, and capable of increasing ARG1 activity in the liver and decreasing the levels of circulating arginine at the same time in a wild-type mouse model with nano gram dosage. Furthermore, treatment of neonatal Arg1 deficiency mouse model with STX-Arg1 EVs resulted in increased life span, because of an active delivery of ARG1 into the liver and reduced circulating arginine. The results presented here demonstrated the efficacy and safety of our STX-Arg1 EVs and showed tremendous potential as enzyme replacement therapy for the treatment of ARG1-D.

## Results

### ARG1 protein expression on engineered cells and EVs

Using Capricor’s proprietary StealthX™ platform, 293F cells were engineered to express ARG1 protein on the EVs membrane. Because localization of ARG1 protein on EVs membrane could affect delivery and its activity, three different constructs were designed to allow the expression of ARG1 protein as 1) a free form (Arg1-free), 2) on the cell surface by linking to the N-terminal of the tetraspanin CD9 (STX-Arg1-out), or 3) tethered to the C-terminal of the CD9 (STX-Arg1-in) **(Figure 1A)**. ARG1 protein expression was confirmed by flow cytometry (**Figure 1B**) and Jess Western blot (**Figure 1C**). As shown in **Figure 1B**, localization of ARG1 matched the original design (as illustrated in Figure 1A). Enrichment of exosome markers (CD9, CD81) and absence of cellular markers (calnexin, GM130, Hsp60) was confirmed by Jess Western blot (**Suppl. Figure 1**).

**Figure 1.**
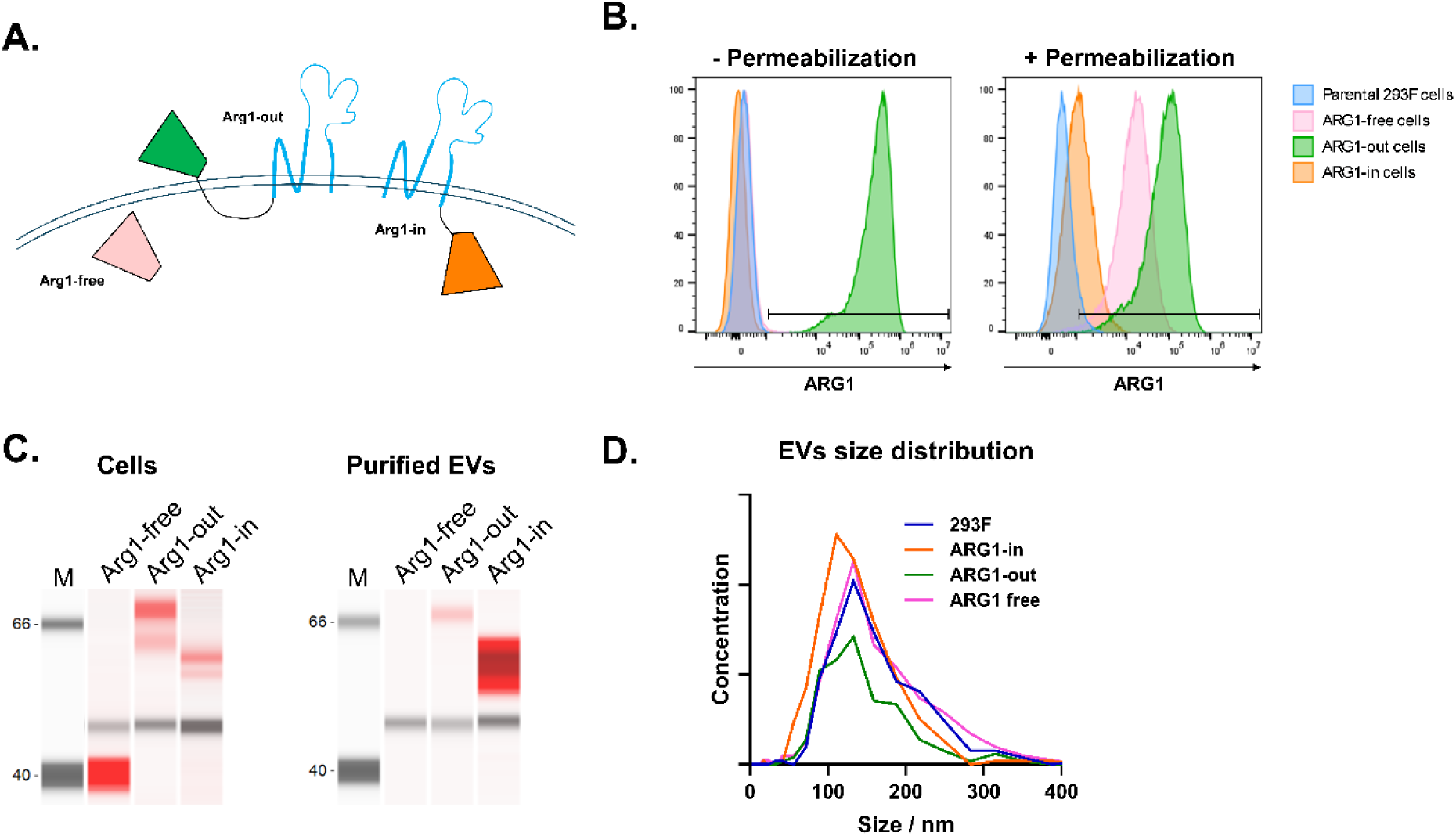
Characterization of StealthX™-engineered cells and EV. **(A)** A scheme showing the design of the expression of ARG1 protein with different anchor designs and orientations on the cell surface (pink: free ARG1; green: STX-ARG1-out; orange: STX-ARG1-in; blue: tetraspanin CD9). **(B)** Flow cytometry analysis of the ARG1 protein expression on different ARG1 cell lines. Parental 293F cells served as a control and no expression of ARG1 protein was detected. Extracellular ARG1 expression was only detected on Arg1-out cells (-permeabilization). ARG1 protein expression on Arg1-free and Arg1-in cells can only be detected after fixation and permeabilization of the cells (+ permeabilization) (pink: free ARG1; green: STX-Arg1-out; orange: STX-Arg1-in; blue: parental cell line) **(C)** ARG1 expressions on cells and EVs were confirmed by Simple Western blotting. Near infrared red: ARG1 (STX-Arg1 in: ∼60 kDa; STX-Arg1-out: ∼70 kDa; free Arg1: ∼ 40 kDa). Chemiluminescence: actin, used as loading control ∼50 kDa. **(D)** Size distribution of 293F and STX-Arg1 EVs by ZetaView nanoparticle tracking analysis (NTA).

Next, EVs carrying ARG1 protein (STX-Arg1) were produced and purified from the culture supernatants. The concentration of the purified EVs ranged between 0.96E12 – 3.0E12 particles/ml with an average diameter of 120.3 – 129.5 nm and a polydispersity index (PDI) of 0.133 – 0.151 **(Figure 1D**, **Table 1)**. Enrichment of ARG1 protein in the EVs preparations was confirmed by Jess Simple Western analysis for both STX-Arg1-in and STX-Arg1-out, but not Arg1-free EVs **(Figure 1C)**. Furthermore, the ARG1 protein carried by 1E11 EV/ml of -out and Arg1-in EVs were 7.0 ± 0.5 and 8.7 ± 0.2 ng/ml with the arginase activities at 51.1 ± 1.3 and 226.4 ± 5.7 U/L, respectively **(Table 1)**. Arg1-free EVs, on the other hand, showed minimal detection of ARG1 protein by Simple Western Protein Analyses **(Figure 1C, right panel)** with 3.7 ± 0.4 ng/1E11/ml EVs and 6.0 ± 0.0 U/L arginase activity **(Table 1)**. Due to the low arginase activities from the Arg1-free EVs, only Arg1-out and Arg1-in were further studied for the in vitro delivery of ARG1 protein.

**Table 1.** Summary of STX-Arg1 EVs characteristics.

|  | EVs per 1E+06 cells<sup>d</sup> | Concentration<br>(EVs/mL) | Size (nm) | PDI | Total protein<br>concentration<sup>a</sup><br>(mg/mL) | ARG1<br>concentration<sup>b</sup><br>(ng/mL) | Arginase<br>activity<sup>c</sup><br>(U/L) |
| --- | --- | --- | --- | --- | --- | --- | --- |
| Arg1-free | 1.19E+10 | 0.96E12 | 129.5 ± 49.8 | 0.148 | 0.027 | 3.7 ± 0.4 | 6.0 ± 0.0 |
| Arg1-out | 8.33E+09 ± 3.17E+09 | 2.20E12 | 129.4 ± 47.2 | 0.133 | 0.026 | 7.0 ± 0.5 | 51.1 ± 1.3 |
| Arg1-in | 1.15E+10 ± 5.21E+09 | 3.03E12 | 120.3. ± 46.7 | 0.151 | 0.025 | 8.7 ± 0.2 | 226.4 ± 5.7 |

<sup>a</sup>: Total protein concentrations were shown in 1E11 EVs/ml
<sup>b</sup>: ARG1 concentrations were shown in 1E11 EVs/ml
<sup>c</sup>: ARG1 activities were shown in 1E11 EVs/ml
<sup>d</sup>: For comparison, 293F cells yield 6.02E+09±1.47E+09 EVs

### In vitro delivery of ARG1 by Arg1-in EVs

To evaluate the enzyme activity of STX-Arg1 EVs and test their ability to deliver functional ARG1 protein into the cells, EVs carrying ARG1 anchored on their membrane (STX-Arg1-out and STX-Arg1-in) were compared to the recombinant human ARG1 protein (rHuArg1) for catalyzation of arginine and intracellular delivery of arginase activities. A hepatocellular carcinoma cell line, HepG2 cells, was treated with 1E11 Arg1-in EVs (carrying 8.7 ng ARG1 protein; 1X), 1E11 (0.5X) or 1.96E11 (1X) Arg-out EVs (carrying 4.5 or 8.7 ng ARG1 protein), or 8.7 ng rHuARG1 for 6, 24, and 48 hours (hr). Urea concentrations in the culture supernatants and arginase activities in the cell lysates were measured. The results showed that both urea concentration **(Figure 2A)** and intracellular arginase activity **(Figure 2B)** were not affected by the treatment of 8.7 ng rHuArg1 protein across the time points studied. In contrast, 1E11 of both STX-Arg1-out and STX-Arg1-in EVs showed a significant increase of urea concentration in the culture supernatants in a time-dependent manner **(Figure 2A)**. 1E11 STX-Arg1-out EVs, which carried ∼50% less ARG1 protein compared with rHuArg1 protein control, showed similar urea production compared to 1E11 STX-Arg1-in EVs. In addition, 1.96 E11 STX-Arg1-out EVs (8.7 ng ARG1 protein) showed significant higher urea production compared to STX-Arg1-in at 6 (p = 0.0103), 24 (p = 0.0407), and 48 (p = 0.0159) hr after treatment, suggesting ARG1 protein carried outside of the EVs can catalyze arginine more efficiently. In contrast, STX-Arg1-out EVs had limited intracellular delivery of ARG1 protein, similarly to the rHuArg1 **(Figure 2B)**. STX-Arg1-out-treated HepG2 cells showed only a limited increase of arginase activity 6 hr after treatment and the arginase activity was undetectable afterward, despite the dose used **(Figure 2B)**. On the other hand, treatment with STX-Arg1-in EVs increased intracellular arginase activities in the HepG2 cells, and the level was higher than both rHuArg1 and the two concentrations of Arg1-out EVs tested. Additionally, after STX-Arg1-in treatment, arginase activities remain elevated 48 h after treatment **(Figure 2B)**. As control, urea production detected in the culture supernatant from the culture media treated with rHuArg1 or STX-Arg1 EVs in the absence of cells was tested and reported in **Suppl. Fig. 2A,** showing bioactivity. To exclude the possibility that EVs are interacting with the plasma membrane of HepG2 cells and triggering intracellular signals to promote specific functions, HepG2 cells were treated with either 293F parental EVs, STX-Arg1-in EVs or rHuArg1. As shown in **Suppl. Figure 2B**, 293F EVs did not trigger any Arg1 activities in cells, confirming that the resulting effect is due to the Arg1 delivered by STX engineered EVs. Overall, the data showed that although ARG1 protein displayed on the surface of EVs (Arg1-out) were more potent in catalyzing arginine, only ARG1 protein encapsulated inside of the EVs (Arg1-in) could deliver ARG1 protein into the cells and maintain its enzymatic activity for longer period of time.

**Figure 2.**
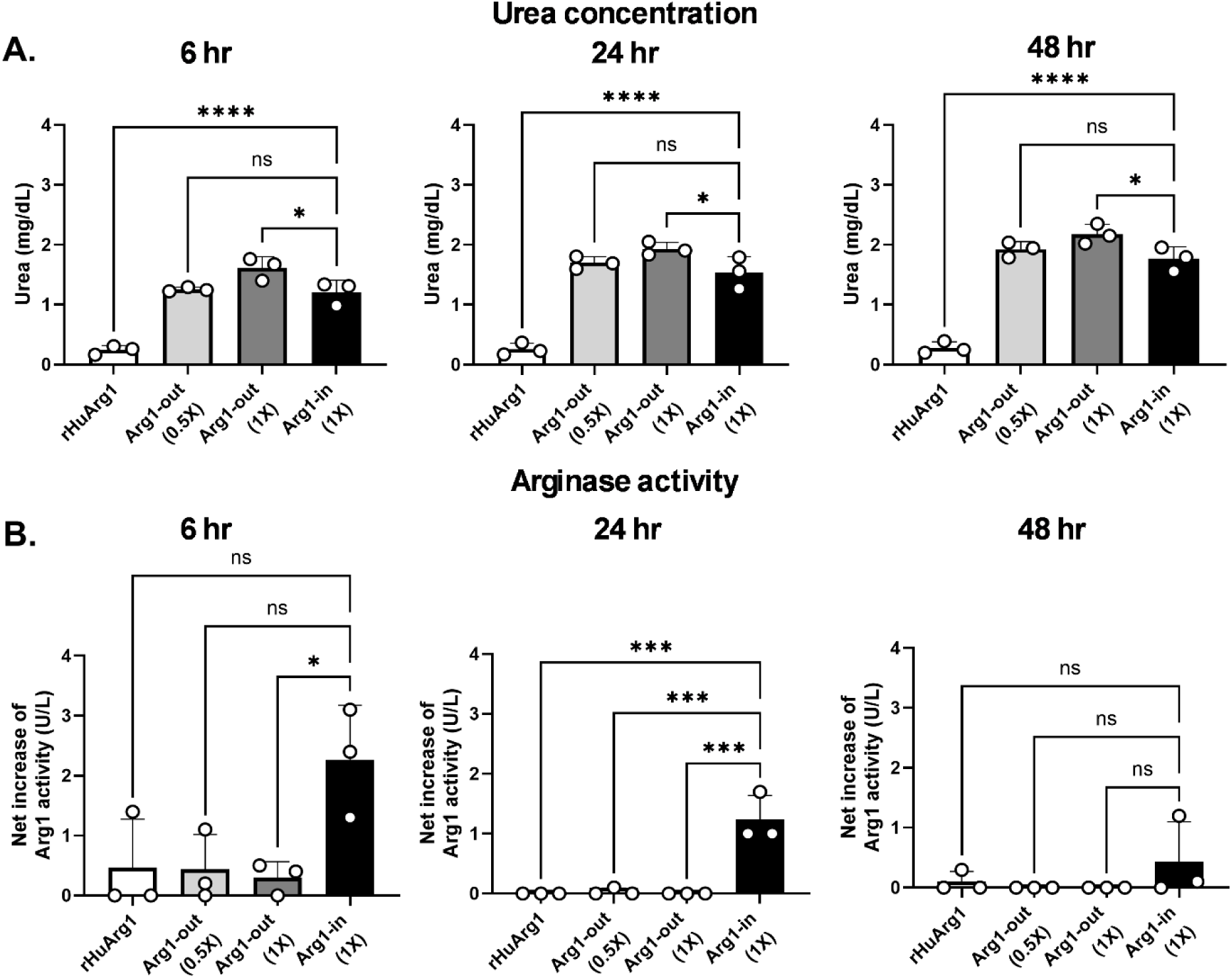
Evaluation of intracellular ARG1 delivery by ARG1 EVs in vitro. HepG2 cells were treated with 1X (1E11; 8.7 ng ARG1) Arg1-in EV, 0.5X (1E11; 4.5 ng ARG1) or 1X (1.96E11; 8.7 ng ARG1) Arg1-out EV, or 8.7 ng rHuArg1 protein for 6, 24, and 48 hr. Culture supernatants and cell lysates were collected separately to evaluate the enzymatic activity of ARG1. **(A)** Urea concentrations in the culture supernatant. Urea productions in both Arg1-in and Arg1-out EV-treated cell cultures were time-dependent. ARG1 carried by EVs was more efficient in converting arginine into urea compared to the same concentration of rHuArg1. **(B)** Arginase activities in the cell lysates. Unlike rHuArg1 and Arg1-out EVs failed to achieve intracellular delivery of ARG1, increased arginase activities can be readily detected in the HepG2 cells 6 h after Arg1-in EV treatment and gradually decreased overtime. Data was analyzed using an ordinary one-way ANOVA with Tukey’s multiple comparison test. A P value < 0.05 was considered to be statistically significant. ns: not significant. *: P < 0.05. **: P < 0.01. ***: P < 0.001. ****: P < 0.0001.

### Potency of Arg1-in EVs

Since STX-Arg1-in EVs demonstrated a better intracellular delivery of functional ARG1 protein, additional studies were performed to elucidate the potency of ARG1 EVs. First, a dose response study was performed to identify the optimal dose (EVs amount, ARG1 concentration) for STX-Arg1-EVs: as shown in **Suppl Figure 3**, a dose of 1E11 EVs resulted in the higher ARG1 activity and urea production, therefore it was chosen for subsequent studies. Next, STX-Arg1-in EVs was compared to increasing doses of rHuArg1: urea levels and intracellular delivery of ARG1 protein were used as endpoint to assess their respective potency. HepG2 cells were treated with 1E11 STX-Arg1-in EVs (8.7 ng ARG1), or 8.7, 50, 250, or 1250 ng of rHuARG1, and the urea production and arginase activities were monitored. In consistent with previous data, urea concentrations in the culture supernatants from HepG2 cells treated with STX-ARG1-in EVs were detected 6 hr post treatment and gradually increased afterward **(Figure 3A)**. Similar trends of the increase in urea concentration were also observed in the culture supernatant from HepG2 cells treated with rHuArg1 at 1250, 250, and 50 ng/well but not 8.7 ng/well **(Figure 3A)**. STX-Arg1-in EVs produced similar levels of urea compared to 1250 ng rHuArg1, and significantly higher than the 8.7 ng (P < 0.0001), 50 ng (P < 0.0001), and 250 ng (P < 0.05 for 6 and 24 hr timepoints) of rHuArg1. Intracellular arginase activity was similarly affected: STX-Arg1-in EVs-treated HepG2 cells showed the highest increase of arginase activities 6 hr after treatment, with decline by 48 hr timepoint **(Figure 3B)**. In contrast, rHuArg1 showed no increase of the arginase activity across timepoints regardless of the concentrations **(Figure 3B),** indicating that STX-Arg1-in had better cell uptake compared with rHuArg1. Overall, STX-Arg1-in EVs outperformed rHuArg1: efficient intracellular delivery with higher enzymatic activity at lower dose.

**Figure 3.**
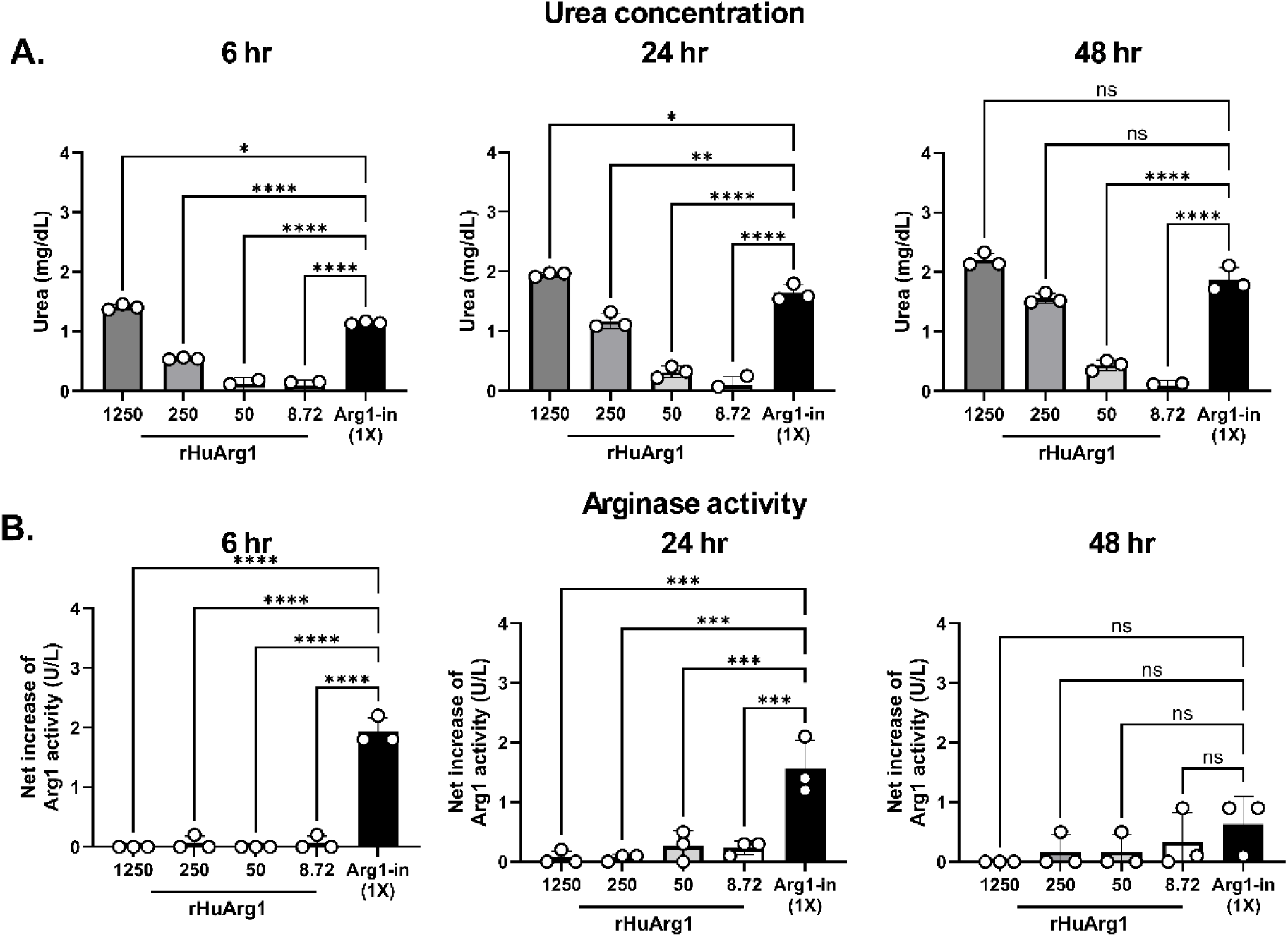
Potency of Arg1-in EVs compared to rHuArg1. HepG2 cells were treated with 1E11 (8.7 ng ARG1) Arg1-in EVs or with a scalar dose of rHuArg1 (8.7, 50, 250, 1250 ng), and culture supernatants and cell lysates were harvested separately 6, 24, and 48 hr after treatment to evaluate the potency of the Arg1-in EVs. **(A)** Urea concentrations in the culture supernatant. Urea concentrations from the cell cultures treated with Arg1-in EVs carrying 8.7 ng ARG1 were comparable to the cells treated with 1250 ng of rHuArg1 and were significantly higher than the ones from rHuArg1 at 8.7, 50 and 250 ng-treated cells across the time points tested. **(B)** Arginase activities in the cell lysates. Arg1-in EVs treatment significantly increased the intracellular arginase activity in HepG2 cells. Arginase activity in rHuArg1-treated cells remained at background level despite the highest concentration tested was 143.7-fold higher than the Arg1-in EVs. Data was analyzed using an ordinary one-way ANOVA with Tukey’s multiple comparison test. A P value < 0.05 was considered to be statistically significant. ns: not significant. *: P < 0.05. **: P < 0.01. ***: P < 0.001. ****: P < 0.0001.

### Biodistribution of STX-Arg1 EVs in vivo

To address the biodistribution of STX-Arg1 EVs, wild type mice received labeled STX-Arg1-in EVs, and their distribution was analyzed on IVIS imager (PerkinElmer) (**Figure 4A-C**). The distribution pattern of the STX-Arg1-in EVs resembled the parental 293F EVs. After injection, STX-Arg1-in EVs, similarly to the parental 293F EVs, rapidly distributed to the liver, spleen and kidney. After 24 hr, kidney signal increased, indicating clearance. One week (wk) after injection, STX EVs signal was cleared from the liver, and minimal signal was recorded in the kidney and bladder, indicating complete clearance **(Figure 4 C, Suppl. Figure 4)**. The data showed that the biodistribution of STX-Arg1-in EVs was not altered by the expression of the ARG1 protein and aligned with published data on EVs distribution (24).

**Figure 4.**
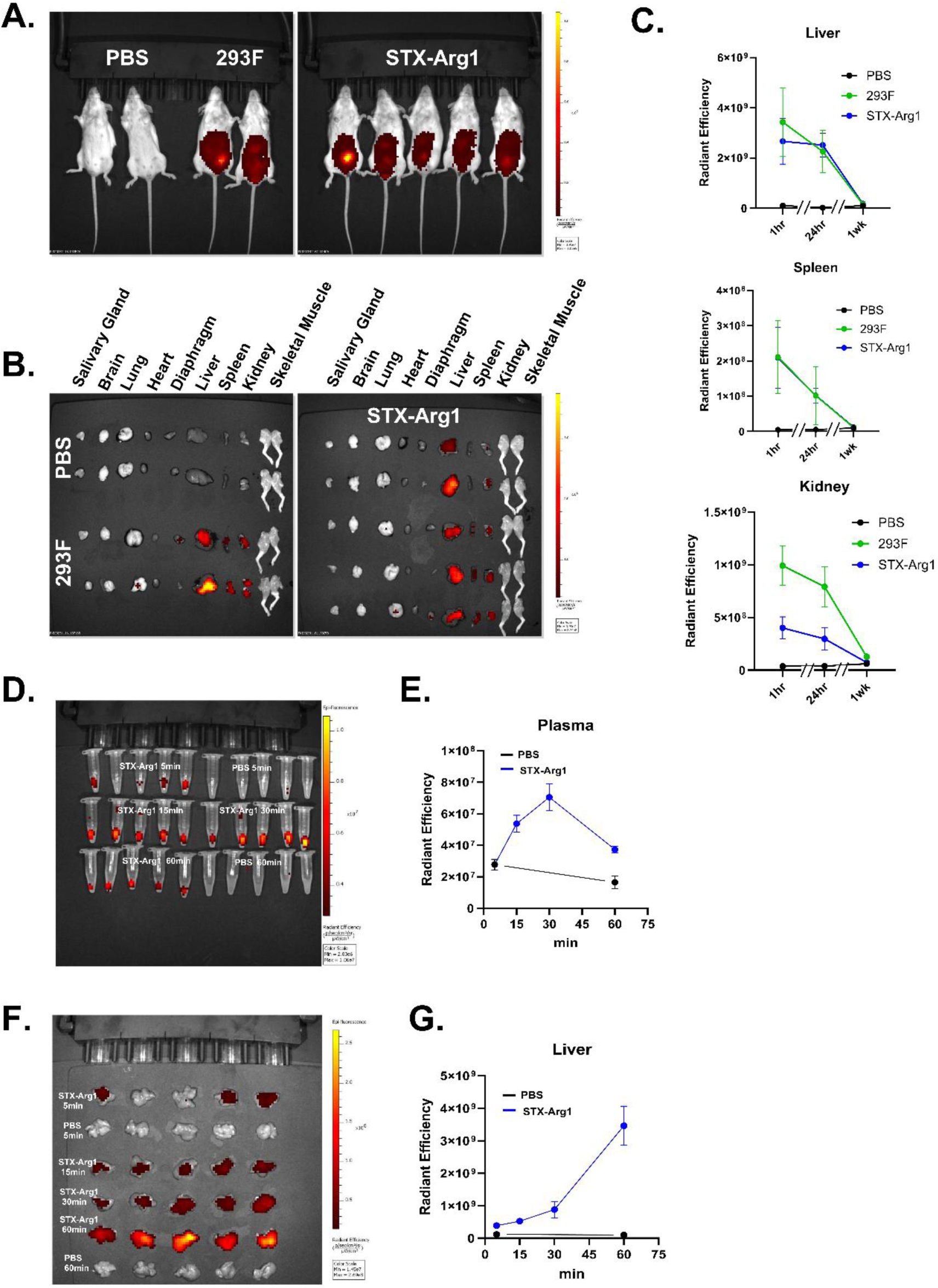
Biodistribution of STX-Arg1 EVs in vivo. Biodistribution of STX-Arg1-in EVs was evaluated in vivo by IVIS imaging. **(A)** Representative IVIS image of PBS control mice, 293F EVs, STX-Arg1-in EVs of injected mice. **(B)** Representative IVIS image of peripheral tissues collected from PBS control, 293F EVs and STX-Arg1-in EVs injected mice. **(C)** Quantification of EVs in tissues at 1hr, 24hr and 1 week after injection. **(D)** IVIS image of plasma collected from PBS and STX-Arg1-in EVs injected mice, at different time points. (**E)** Quantification of STX-Arg1-in EV signal in plasma (from E). Black line is PBS injected mice, blue line is STX-Arg1-in EVs mice. **(F)** IVIS image of livers collected from PBS and STX-Arg1-in EVs injected mice, at different time points. **(G)** Quantification of STX-Arg1-in EV signal in plasma (from G). Black line is PBS injected mice, blue line is STX-Arg1-in EVs mice. N=5 mice per experimental treatment.

To evaluate pharmacokinetic of STX-Arg1-in EVs, blood was collected at 1hr and 24 hr to evaluate the blood retention of ARG1 EVs. As shown in **Figure 4**, STX-Arg1-in EVs were detected in blood 1 hr after injection, mainly in plasma. No signal was detected in the cellular fraction. The EVs-associated signal was completely cleared from circulation by 24 hr post-injection (**Suppl. Figure 5**).

In a separate experiment, STX-Arg1-in EVs pharmacodynamic was evaluated at shorter timepoints (**Figure 4D-G**). Blood and liver were collected from mice receiving either PBS or labeled STX-Arg1-in EVs at 5, 15, 30, 60 minutes (min) post-injection. STX-Arg1-in EV-associated signal was quantified and analyzed. As shown in **Figure 4D-G**, after intraperitoneal injection, STX-Arg1-in EVs rapidly distributed to the circulation, with a peak at 30 min. The STX-Arg1-in EVs were removed from the circulation and distributed to the liver, in the following 30 min (**Figure 4D-G**). This data aligns with the PD/PK of other EVs (24).

### In vivo toxicity using wild-type mice

In vivo functionality and toxicity of STX-Arg1-in EVs was first addressed in wild-type mice. Age matched 8-10 wk old BALB/C mice were assigned to either control group (receiving vehicle PBS) or STX-Arg1-in EVs at max calculated dose allowed by manufacture yield of 0.012 mg/kg. Two dosing schedules were implemented: one cohort received one i.p. injection per week, and one cohort received two i.p. injections per week (**Figure 5A**). Weight was monitored weekly, and regardless of the dosing schedule, no significant changes in weight were recorded across treatments (**Figure 5B**). Levels of circulating arginine were quantified by ELISA after terminal blood collection. Lower levels of arginine were observed in the blood of mice receiving one i.p. injections of STX-Arg1-in EVs per week compared to PBS (**Figure 5C**). A trend of decrease in arginine concentrations, although not statistically significant, was observed in mice receiving 2 i.p. injections of STX-Arg1 per week (**Figure 5C**). No changes in AST levels were observed in either dosing group (**Figure 5D).** Peripheral tissues (see Methods) were collected for histology evaluation. At the more frequent dosing schedule, few microscopic alterations were observed, which included minimal to mild leukocyte infiltrates around an airway across all treatments in the lung, 1 µm or smaller cytoplasmic indistinct vacuoles in the liver, minimal to no lymphoid hyperplasia, across all treatments in the spleen and mildly dilated renal tubules in the kidney (**Figure 5E)**. All findings were defined as not adverse, suggesting a safe profile of STX-Arg1-in EVs.

**Figure 5.**
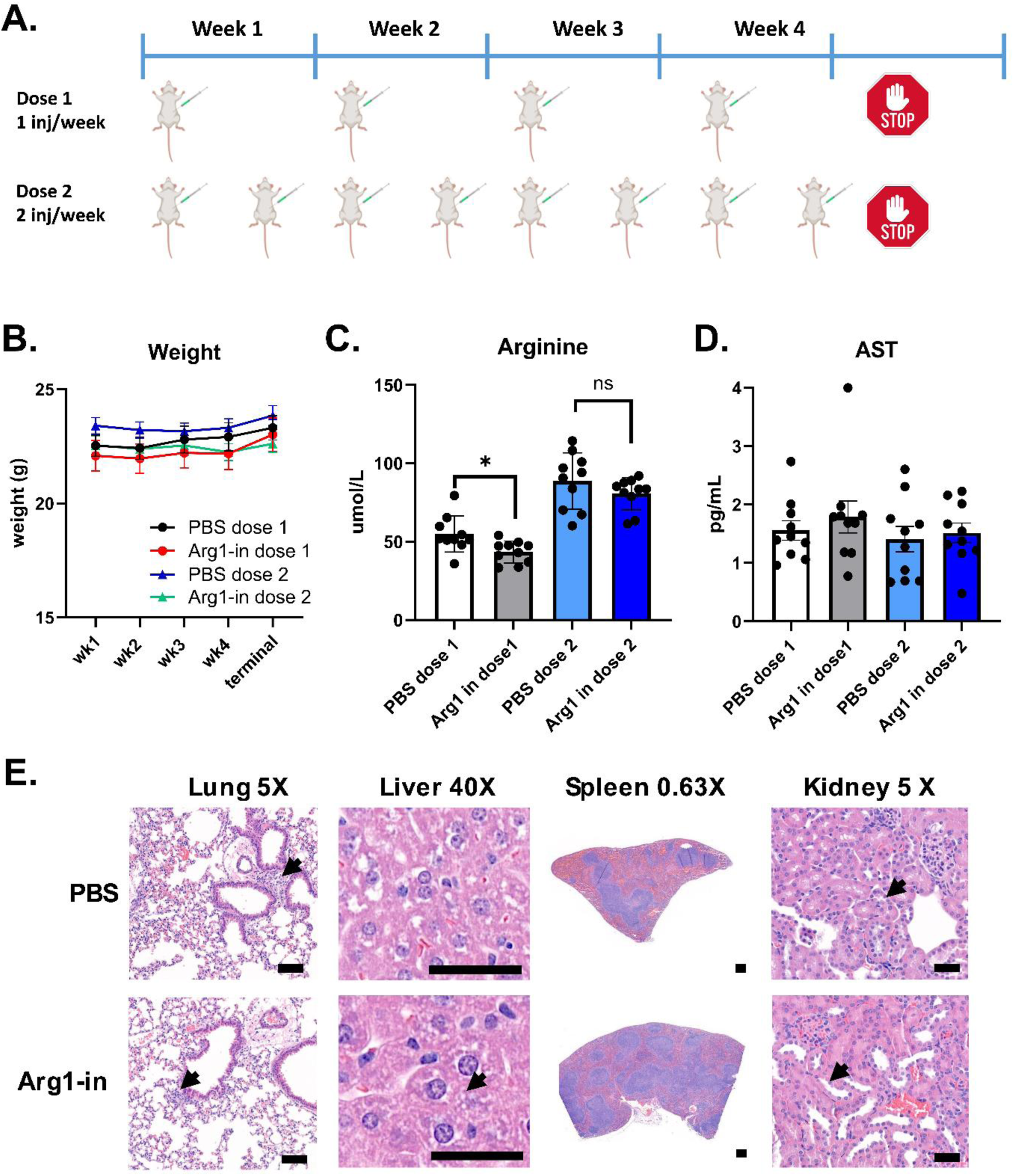
In vivo toxicity of STX-Arg1-in EVs injection in wild-type mice. **(A)** Schematic representation of the dosing schedule. **(B)** Weight was monitored weekly. **(C-D)** Percentage of change of circulating arginine in mice receiving STX-Arg1-in EVs, compared to PBS group. **(E)** Representative images of H&E-stained tissues from PBS-and STX-Arg1-in-treated mice. Arrows indicate alterations. Scale bar: 50µm for lung, Liver and kidney; 100µm for spleen. N = 10 mice per experimental group. 2-tailed t-test, ns: not significant, *: p < 0.05 N=10 mice per experimental treatment.

### In vivo delivery of ARG1 protein and efficacy in Arg1 knockout mouse model

To address the therapeutic potential of STX-Arg1-in EVs, Arg1 knock-out mouse model (Arg1^-/-^) neonatal mice received i.p. injection of STX-Arg1-in at a maximum dose of 0.03 mg/kg. Repeated dosing resulted in improved survival in Arg1^-/-^ neonatal pups with arginase deficiency (**Figure 6A**) and weight gain comparable to littermates **(Figure 6 B)**: while Arg1^-/-^ pups receiving vehicle PBS died by day 14 (similarly to what reported in the literature, (7)), 45 % of the Arg1^-/-^ pups undergoing STX-Arg1-in EVs therapy survived day 14, while 25% survived day 18. A two-way ANOVA analysis showed a p=0.0022, suggesting the effectiveness of the treatment. The positive effect on lifespan was to be attributed to the delivery of ARG1 by EVs to the liver cells.

**Figure 6.**
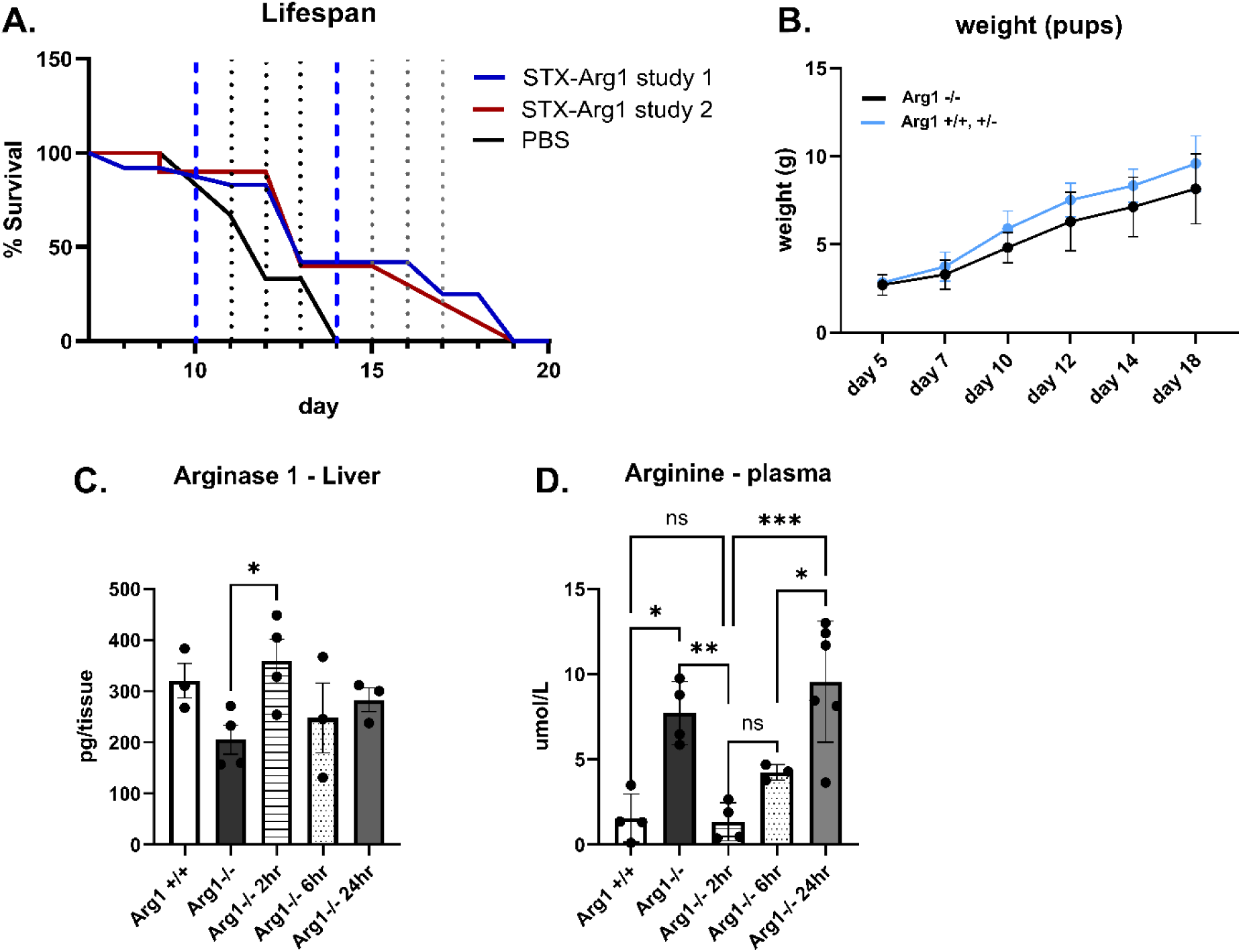
Efficacy of STX-Arg1-in EVs therapy in Arg1^-/-^ mice. **(A)** Lifespan analysis. Blue segmented lines represent the window of mortality observed in Arg1^-/-^ mice. Black dots lines identify the single days. 2-way ANOVA showed a p=0002 for treatment effect. **(B)** Weight assessment through treatment. (**C**) ARG1 levels in livers of naïve wild type Arg1^+/+^ and knock-out (Arg1^-/-^) mice, and after treatment in Arg1^-/-^ mice. **(D)** Arginine levels in livers of naïve wild type (Arg1^+/+^) and knock-out (Arg1^-/-^) mice, and after treatment in Arg1^-/-^ mice. N = 4 Arg1^-/-^ mice assigned to PBS treatment. N = 22 Arg1^-/-^ mice assigned to ARG1 treatment. One-way ANOVA adjusted for multiple comparisons, ns= not significant, *: p < 0.05, **: p < 0.01, ***: p < 0.005.

A time-course study was performed on the Arg1^-/-^ mice to address the effect of STX-Arg-in on lifespan in this model. Arg1^-/-^ pups received the STX-Arg1-in injection and both blood and liver were collected 2, 6 and 24 hr after the injection. As shown in **Figure 6C**, STX-Arg1-in EVs injection restored liver ARG1 levels in the Arg1^-/-^ pups by 2 hr, up to the levels of wild-type littermates, but it was rapidly cleared afterwards starting at 6 hr. Consequently, arginine levels in circulation rapidly declined at 2 hr in Arg1^-/-^ undergoing STX-Arg1-in treatment, reaching levels observed in wild-type littermates, and steadily returning to baseline levels by 24 hr (**Figure 6D**). Arg1^-/-^ mice gained weight throughout the study **(Figure 6B)**, but lethargic phenotype was observed at later life stages, possibly due to hyperammonemia, which is the leading cause of death in this model (7).

## Discussions

Currently, the treatment for ARG1-D mainly rely on the restriction of dietary intake of protein, arginine-free essential amino acid supplement, and nitrogen-scavenging drugs, such as glycerol phenylbutyrate or sodium phenylbutyrate, as needed. Although numerous strategies, including AAV-based gene therapy (8–11), lipid nanoparticle-based mRNA therapy (12), recombinant ARG1 protein (7), hepatocyte transplantation (25), and stem cell-based therapy (25–27), have been evaluated for the treatment for ARG1-D for the past two decades, minimal progress was made to the clinical. Some approaches were shown to be with limited efficacy (7) and the rest were hindered by various challenges, including pre-existing immune responses against gene therapy vectors and immune-mediated toxicities after administration of AAV vectors (28, 29), genotoxicity of AAV-based therapies (30, 31), inability of intracellular delivery of hepatocytes (7), and safety and adverse effects of synthetic lipid nanocarriers in pediatric cohorts (32). Thus far, only one disease-specific drug, a cobalt-substituted pegylated recombinant human ARG1 targeting hyperargininemia, was approved until recently in the EU and UK (https://www.ema.europa.eu/). It is worth noting that while hyperammonemia is typically not present, the ARG1-D diagnosis is increasingly made by expanded newborn screening. Therapeutic drugs that can directly improve the deficit of ARG1 protein in ARG1-D remain direly needed.

In this study, we developed an EV-based enzyme replacement therapy that successfully prolonged the lifespan of a lethal neonatal Arg1^-/-^ mouse model. By generating EVs carrying enzymatic active ARG1 protein on the EV surface or inside of the EV lumen, we showed that the capability in catabolizing arginine by EVs carrying nanogram of ARG1 were similar to microgram level of fee rHuArg1. We further demonstrated intracellular delivery of functional ARG1 protein by EVs in vitro. In vivo studies demonstrated that STX-Arg1-in EVs were not toxic. Most importantly, the lifespan of neonatal Arg1^-/-^ mouse model was extended by the treatment of STX-Arg1-in EV due to the successful delivery of ARG1 enzyme to the liver, retaining its enzymatic activities and reducing arginine level in circulation.

EVs were chosen as the vehicles for the cargo delivery of engineered human ARG1 protein. EVs are cell-derived lipid nanoparticles that serve as a messenger for communication between cells. Due to the nature of the EVs, these cell-derived nanovesicles are considered to be safe, have low immunogenicity, and high biocompatibility (16, 21). Decades long application of blood transfusion, which contains high concentrations of EVs, further strengthening the safety of this approach. In our previous studies using StealthX™ EV platform, we demonstrated a high safety profile of the STX EVs (33, 34). This was further supported by the current study: no adverse effect, toxicity, or pathological histopathological changes were found in the wild-type mice receiving STX-Arg1-in EVs **(Figure 5)**. The high safety profile allowed us to administrate multiple doses for therapeutic purposes.

The goal of the present study was to develop a novel enzyme replacement therapy for ARG1-D using EV-based approaches to reduce the amount of protein needed for therapeutic efficacy and improve delivery to the liver for the treatment of ARG1-D. As anticipated, the localization of the ARG1 protein on EV membrane affected its functionality. Using our StealthX™ EV-engineering platform, ARG1 protein was loaded to the EVs through an EV-specific anchor (STX-Arg1-out and STX-Arg1-in) or an anchor-free (Arg1-free) system (**Figure 1A**). In alignment with previous data, EV-specific anchor provided a more efficient loading of ARG1 protein to the EVs, with STX-Arg1-out and STX-Arg1-in EVs showing higher arginase activities compared to the STX-Arg1-free EVs **(Figure 1D)**. To gain a better insight into the potential efficacy, in vitro delivery of ARG1 protein by ARG1 EVs was studied. Our data showed that nanogram-levels of ARG1 protein delivered by STX-Arg1-out and STX-Arg1-in EVs had comparable enzyme activities in catalyzing arginine to microgram-level of rHuArg1 **(Figures 2A and 3A)**. As urea cycle takes place in the hepatocytes in the liver and catabolizes the toxic ammonia into urea (2), expression of functional ARG1 in the hepatocytes has been suggested to be important to resolve hyperammonemia and prolong survival of Arg1 knockout mouse models (7–9). Our data showed that STX-Arg1-in EVs, which have the engineered ARG1 protein encapsulated inside of the EVs, could deliver ARG1 into the cells and retain its function, which was not achievable by either STX-Arg1-out EVs **(Figure 2B)**, or rHuArg1 **(Figure 3B)**. This difference in activity between the two constructs (STX-Arg1-in and STX-Arg1-out) might be due to the localization of Arg1 protein in respect to the CD9+linker: the linker might be affecting its enzymatic activity. Moreover, it could be also possible that localization of the enzyme in the extra vesicle space exhausted its enzymatic activity faster, but could still be useful for clearance of the circulating arginine.

Enrichment of STX-Arg1-in EVs in the liver **(Figures 4 and 5, Supplementary Figure 4)** further emphasized the potential of our strategies for the treatment of ARG1-D. The lethal phenotype of the animal models for the ARG1-D (Arg1^-/-^ mouse) makes the development of therapeutics challenging. Unlike human ARG1-D, that the disease rarely leads to death by the symptoms due to lack of ARG1 activity directly, both neonatal/congenital and inducible/conditional Arg1-knockout mouse models were lethal (7, 27), with a lifespan of 14 days from birth or after induction. The main cause of death of Arg1-knockout mouse models were considered to be hyperammonemia (7, 27), which was less frequent in human ARG1-D (3). In our study, the therapeutic efficacy of the STX-Arg1-in EVs was evaluated on a neonatal Arg1^-/-^ mouse model, B6.129-Arg1tm1Rki/J (The Jackson Laboratory, Strain # 007741). STX-Arg1-in rapidly delivered ARG1to the liver, which resulted in rapid decrease of circulating plasma arginine. A fast clearance was observed, with decline 6 hr after injection **(Figure 6)**. More importantly, our data showed that i.p. administration of STX-ARG1-in EVs (at a dose of 0.03 mg/kg of ARG1Arg1) weekly or every 48 hours was sufficient to extend the lifespan of the Arg1^-/-^ mice by 45 % over day 14, and 25% over day 18 (**Figure 5**). This result was not achieved by cobalt-substituted pegylated recombinant human ARG1 (7): the authors suggested that the lack of improvement of survival and control of hyperammonemia may be caused by the inability of pegylated recombinant human ARG1 entering the liver although the treatment was demonstrated to be efficient in controlling arginine concentrations in the circulation. The capability of STX-Arg1-in EVs in controlling blood arginine level in conjunction with extended lifespan of the Arg1^-/-^ mice strengthens the potential of this strategy as a disease-specific approach in treating ARG1-D.

We are aware of the limitations of the present study. While an improvement of the lifespan was observed after STX-Arg1-in treatment, no long-term survival was achieved. Longer lifespan was achieved by AAV- and LNP-mRNA-based therapies, confirming the need of intracellular delivery of the ARG1 and constant supply of the enzyme. Further studies with increased amount of ARG1 and new formulations to increase ARG1 half-life are needed to improve the beneficial effects of STX-Arg1-in EV therapy. Additionally, a combination of STX-Arg1-in and STX-Arg1-out could be formulated as a cocktail therapy to combine the potent STX-Arg1-out activity for controlling the peripheral arginine levels, with the intracellular delivery capability of STX-Arg1-in. Both STX-Arg1-out (data not shown) and STX-Arg1-in **(Figures 4 and 5)** were tested in vivo in wild-type mice. Both EVs types were biologically active, with no toxic effects, despite the limiting data available.

Lastly, the manufacturing of EV based therapeutics must be considered. Many approaches have been proposed for EVs manufacturing (35): at Capricor, a scalable process has been developed, using tangential filtration and size exclusion chromatography. This approach has been successfully used for the production of clinical doses for vaccine applications and shows scalable capabilities to support therapeutical applications.

Altogether, our data demonstrated the capability of the STX-Arg1-in EVs generated by our StealthX™ platform in intracellular delivery of enzymatic active ARG1 protein, which was further translated to an increased arginase activity in the liver, a relief of hyperargininemia, and most importantly, an extended survival time of a lethal neonatal Arg1-deficiency mouse model. With the safety of the EV-based therapeutics, STX-Arg1-in EVs showed a potential in providing a better therapeutic effect for the patients suffering from ARG1-D.

## Methods

### Cell lines

Human embryonic kidney 293 T cells (293T) were purchased from ATCC (CRL-3216) and were cultured using Dulbecco’s Modified Eagle Medium (DMEM), high glucose, Glutamax™ containing 10% fetal bovine serum at 37℃ with 5% CO_2_. FreeStyle™ 293F cells (Gibco) were purchased from ThermoFisher Scientific. 293F cells were served as a parental cell line to generate stable cell lines expressing human ARG1 protein. Parental 293F cells and the engineered 293F cells were cultured in a Multitron incubator (Infors HT) at 37℃, 80% humidified atmosphere with 8% CO_2_ on an orbital shaker platform rotating at 110 rpm.

### Lentiviral vectors

Lentiviral vectors for expression of human ARG1 (NCBI Reference Sequence: NM_001244438.2) were designed and codon-optimized for the synthesis from Genscript together with the two lentiviral packaging plasmids, pMD2.G and psPAX2. Three different designs were utilized: 1) overexpression of ARG1 (Arg1-free), 2) extracellular display of ARG1 (Arg1-out) by linking ARG1 with CD9 with a synthetic transmembrane linker at the N-terminal of the ARG1 sequence, and 3) intracellular packaging of ARG1 (Arg1-in) by linking ARG1 to the C-terminal of the CD9. Production of lentiviral particles for transduction of the cells was performed as previously ((33, 34). Briefly, lentiviral particles carrying the gene of interest were generated by transfecting 293T cells with pMD2.G (Genescript), psPAX2 (Genescript) and ARG1 expressing vectors (Genscript) at a ratio of 5:5:1 using Lipofectamine 3000 according to the manufacture’s instruction. Lentiviral particles were collected at 48 hours after transfection and used for the transduction of 293F cells.

### ARG1 EV production

Suspension Arg1-free, Arg1-out, and Arg1-in cells were cultured in FreeStyle 293 Expression media (Chemical defined, EVs and serum free, ThermoFisher Scientific) in a Multitron incubator (Infors HT) at 37℃ at 80% humidified atmosphere supplied with 8% CO_2_ on an orbital shaker platform. Cell suspension was collected 72hrs after seeding, and the cells and cell debris were removed by centrifugation, while microvesicles and other extracellular vesicles larger than ∼220 nm were removed by vacuum filtration. Clarified supernatants were processed as previously described (33, 34). Briefly, supernatant was subjected to concentrating tangential flow filtration (TFF) on an AKTA Flux s instrument (Cytiva, United States) and then subjected to chromatography on an AKTA Avant 25 (Cytiva, United States).

### Nanoparticle tracking analysis

Size distribution and concentration of purified EVs were determined using ZetaView Nanoparticle Tracking Analysis (Particle Metrix) according to manufacturer instructions. EV samples were diluted in 0.1 µm-filtered 1X PBS (Gibco) to fall within the instrument’s optimal operating range to allow the optimal characterization of the EVs. For reproducibility, the following conditions were applied: 1. Number of positions:11; 2. Resolution: high, 60 frames; 3. sensitivity: 84-87; 4. Shutter: 100; 5. Frame rate: 30 fps; 6. Minimum brightness: 20; 7. Max area: 1000; 8. Min area: 10; 9. Tracelength: 15; 10. Laser: 488; 11: temperature: Room temperature; 12: Replicates: three (3) replicated per sample.

### Flow cytometry

ARG1 protein expressions on the cells were detected by the standard flow cytometry techniques. ARG1 protein positioned outside of the cells and EVs (Arg1-out) can be detected by surface staining, while intra-cellular/intra-vesicle ARG1 (Arg1-in) requires membrane permeabilization to allow its staining/detection. Briefly, for the detection of extracellular ARG1 and CD9, 300,000 cells/well were plated in a 96 wells U bottom plate for staining. Cells were incubated at 4℃ for 20 min with 100 µl eBioscience™ Flow Cytometry Staining Buffer (ThermoFisher Scientific) containing anti-human ARG1 antibodies (Biolegend, clone 14D2C43, PE) protected from light. For intracellular ARG1 (STX-Arg-in) staining, cells were permeabilized and fixed for 20 min at 4℃ using BD CytoFix/CytoPerm buffer (BD Bioscience) followed by incubation at 4℃ for 20 min with 100 µl BD Perm/Wash Buffer (BD bioscience) containing anti-human ARG1 antibodies (Biolegend, clone 14D2C43, PE) protected from light.After staining, cells were washed and re-suspended in 200 µl eBioscience™ Flow Cytometry Staining Buffer (ThermoFisher Scientific) for sample acquisition using CytoFlex S flow cytometer (Beckman Coulter) and the data was analyzed using FlowJo V10 (Becton, Dickinson and Company). Gating Strategy is available in **Suppl. Figure 6**.

### Simple Western Jess automated protein analysis

Detections of ARG1 protein in cells and EVs were carried out using Protein Simple’s Jess capillary protein detection system. Cell and EV samples were lysed in RIPA buffer (ThermoFisher Scientific) supplemented with protease/phosphatase inhibitor (ThermoFisher Scientific), quantified using the BCA assay (ThermoFisher Scientific) and run for detection. To detect engineered ARG1 protein, the separation module 12–230 kDa was used following manufacturers protocol. Briefly, 1 µg of sample and protein standard were run in each capillary, probed with primary antibodies followed by secondary antibodies provided in Jess kits (HRP/IR). Primary antibody used are as following: rabbit anti-human ARG1 (ThermoFisher Scientific, clone 24H4L3, 1:100 dilution), mouse anti-human actin (Novus Biologicals, clone AC-15, 1:20 dilution), rabbit anti-human CD9 (Cell Signaling, clone D9O1A, 1:100 dilution), rabbit anti-human calnexin (Novus Bio, 1:100 dilution), mouse anti-human CD81 (Novus Bio, 1:10 dilution), rabbit anti-HSP60 (R&D system, 1:100 dilution), rabbit anti-human GM130 (R&D system, 1:10 dilution).

### Quantification of recombinant ARG1 protein by ELISA

Human ARG1 protein carried on the ARG1 EVs were quantified by an enzyme-linked immunosorbent assay (ELISA) using precoated ELISA plates (Invitrogen) following the manufacturer’s instructions. Briefly, EV samples were lysed with RIPA buffer (ThermoFisher Scientific) supplemented with protease/phosphatase inhibitor (ThermoFisher Scientific) at 1:1 ratio for 30 min on ice. 1E11/ml or 1E10/ml of lysed EVs were plated to antigen-coated wells together with a biotinylated detecting antibody and incubated at room temperature for 2 hr on an orbital shaker (200 rpm). After the incubation, the lysates were removed and the wells were washed four times with 1X Wash Buffer, and the Streptavidin-HRP were added into the wells and incubated for 1 hr on an orbital shaker (200 rpm). Wells were washed four times after the incubation and TMP Substrate were added into the wells to allow the color to develop. The reactions were then stopped with Stop Solution once the color was developed and absorbance at 450 nm and 620 nm were recorded using a BioTeck Gen5 plate reader (Agilent). The concentration of ARG1 carried on ARG1 EVs were calculated based on the standard curve.

### Arginase activities of ARG1 EVs

Arginase activities of the ARG1 EVs were quantified using a colorimetric arginase assay kit (BioAssay System). EV samples were lysed with lysis buffer containing 0.4% Triton X-100 (Millipore Sigma Aldrich) supplemented with protease/phosphatase inhibitor (ThermoFisher Scientific) for 30 min on ice, and 1E11/ml, 1E10/ml, and 1E9/ml of ARG1 EVs were tested following the manufacturer’s instructions. Briefly, Arginine Substrate containing Mn were incubated with the samples for 2 hr at 37°C and the amount of urea produced by the enzymatic activities of arginase were detected using Urea Reagent by incubating for 1 hr at room temperature. After incubation, optical density at 430 nm of the samples were measured using a BioTeck Gen5 plate reader (Agilent) and the arginase activities in the samples were calculated based on the urea standard.

### In vitro delivery of ARG1 protein by ARG1 EVs

To evaluate the capability and efficacy of EVs carrying ARG1 in delivering ARG1 protein into the cells, HepG2 cells were treated with either ARG1 EVs or recombinant human ARG1 (rHuArg1) (ACROBiosystems) for 6, 24, and 48 hr, and the urea concentration in the culture supernatant and arginase activities in the cell lysate were measured. 1E5 cells were seeded in 12 wells plate 1 day prior to the experiment to allow the cells to attach. To test the in vitro delivery of ARG1 by EVs, Arg1-out or Arg1-in EVs were diluted in 1X PBS to desired concentration ranging between 1E12 – 1E11/ml. For the rHuArg1, the protein was reconstituted in distilled water at 0.5 mg/ml and stored at −80°C before the experiment without repeated free/thaw cycle. Similar to ARG1 EVs, rHuArg1 was further diluted to 1.25 – 0.0087 µg/ml using 1X PBS. Cells were washed once with EMEM/10% FBS followed by the treatment of EVs or rHuArg1. Cells treated with 1X PBS served as a negative control. To ensure the culture condition of the cells, the volume of EVs, recombinant protein, and PBS were 10% of the culture media. For each time point, culture supernatant was collected and centrifuged at 1,000 xg at 4°C to remove the debris before testing the urea concentration using a Urea Assay Kit (BioAssay System) by following the manufacturer’s protocols. Cells were kept on ice and washed once with cold 1X PBS followed by lysing with lysis buffer containing 0.4% Triton X-100 (Millipore Sigma Aldrich) supplemented with protease/phosphatase inhibitor (ThermoFisher Scientific) for 30 min on ice. After incubation, cell lysates were centrifuged at 14,000 xg at 4°C and the supernatant were tested for their arginase activities by arginase activity assay (BioAssay System) by following the manufacturer’s instructions.

### Exosome Labeling

EVs were labeled with IVISense 750 MAL Fluorescent Self-Quenching Dye (Revvity Health Sciences). Briefly, 1 ml of 1E12 particle/ml was incubated with 20 µg dye and incubated at 37C for 4 hr. Free, unbound dye was removed using the Zeba Spin Desalting Columns (Thermofisher Scientific) according to the manufacturer’s instructions.

### Animal study – biodistribution in wild-type mice

Studies were conducted according to the guideline of the Institutional Animal Care and Use Committee (IACUC protocol EB17-004-091). Mice were fed ad libitum and sterile water; housed in groups of five at 22°C/30% humidity and light cycles of 0600–1800 hours with standard nesting material; and allowed free movement. To examine the tissue distribution of ARG1 EVs, age matched BALB/c mice (female, 8-10 wk old) received intraperitoneal injection (100 µl) of either 1) PBS, 2) 293F or 3) STX-Arg1-in EVs. A total of 1E11 particles/100ul were injected and blood and tissues (salivary glands, brain, lungs, heart, diaphragm, liver, spleen, kidney, lower limbs) were collected at 1 hr, 24 hr and 1 wk after injection. Full body and tissue imaging were acquired on the IVIS imager (PerkinElmer). Additionally, blood clearance and liver accumulation were evaluated at shorter timepoints (5, 15, 30, 60 min after injection).

### Animal study – toxicity in wild-type mice

Studies were conducted according to the guideline of the Institutional Animal Care and Use Committee (IACUC protocol EB17-004-091). Mice were fed ad libitum and sterile water; housed in groups of five at 22°C/30% humidity and light cycles of 0600–1800 hours with standard nesting material; and allowed free movement. To address activity and evaluate toxicity of STX-Arg1 EVs repeated injections, age matched BALB/c mice (female, 8-10 wk old) received intraperitoneal injection (100 µl) of either 1) PBS, or 2) STX-Arg1-in EVs, at the highest dose allowed by our manufacturing scale (0.012 mg/kg). Mice were divided into two dose groups: 1) Dose 1 received one i.p. injection per week, and 2) Dose 2 received two i.p. injections per week. Weight was monitored weekly. At the end of the fourth week, mice were anesthetized using isoflurane, blood collected from the submandibular vein (in EDTA coated tubes), and peripheral tissues (salivary glands, brain, lungs, heart, diaphragm, liver, spleen, kidney, lower limbs) processed for histological analysis. Blood was further processed for plasma isolation after centrifugation at 4000 rpm for 5 min at 4°C. Plasma was analyzed for arginine levels (LSBio, see below), as indirect measurements of STX-Arg1-in activity. Liver toxicity was additionally assessed by quantification of aspartate aminotransferase (AST) (Abcam) levels in blood.

### Animal study – efficacy in Arg1 knock-out mice

Studies were conducted according to the guideline of the Institutional Animal Care and Use Committee (IACUC protocol EB17-004-091). Mice were fed ad libitum and sterile water; housed in groups of five at 22°C/30% humidity and light cycles of 0600–1800 hours with standard nesting material; and allowed free movement. To evaluate the therapeutic potential of STX-Arg1 EVs, Arg1 knock-out mice (B6.129-Arg1tm1Rki/J, Strain #:007741; RRID: IMSR_JAX:007741) were acquired from Jackson Laboratory. Colony was expanded from cryo-embryos to obtain heterozygous mice for subsequent breedings. Heterozygote mice are viable and fertile, while homozygous Arg1 mutant completely lack hepatic ARG1 activity, exhibit hyperargininemia, severe symptoms of hyperammonemia (including decerebrate posture, lethargy, and high-frequency tremor of the extremities, particularly the tail) and die between 10-14 days after birth (36, 37). Mice received either an i.p. injection of 10 µl/g every 2 or 4 days for a final dose of 30 µg/kg ARG1 as delivered by STX-Arg1-in EVs. Weight gain and growth was monitored across the study and lifespan recorded. Additionally, a time course study was performed to evaluate the distribution of STX-Arg1-in EVs in the liver of knock-out mice and its activity. After injection, blood and liver were collected at 2, 6 and 24 hr after injection. Liver was lysed in RIPA buffer (ThermoFisher Scientific) with 1X Protease inhibitors (ThermoFisher Scientific). ARG1 levels were quantified by human ARG1 ELISA (Invitrogen). Blood was centrifuged at 4500 xg for 10 min and plasma collected for arginine levels quantification by ELISA (LSBio, see below).

### Arginine ELISA

Arginine levels in blood were quantified by competitive ELISA (LSBio) as per manufacturer’s instructions. ELISA was used as indirect measure of the STX-Arg1-in EVs in vivo. Briefly, 50 ul of either standards or samples are added to the wells together to a fixed quantity of biotin-conjugated target antigen. The antigens in the standards or samples compete with the biotin-conjugated antigen to bind to the capture antibody. Unbound antigen is washed away. An Avidin-Horseradish Peroxidase (HRP) conjugate is then added which binds to the biotin. After removal of unbound HRP-conjugate a TMB substrate is then added which reacts with the HRP enzyme resulting in color development. Reaction is topped with sulfuric acid stop solution and the optical density (OD) of the well is measured at a wavelength of 450 nm: the greater the amount of antigen in the sample, the lower the color development and optical density reading.

### Statistics

For in vitro studies of the delivery of ARG1 by EVs, the differences between ARG1 EVs and the rHuArg1 were tested using ordinary one-way analysis of variance (ANOVA) with Turkey’s multiple comparison test with a pooled variance to compare the urea concentration in the culture supernatants and the increase of arginase activities in the cell lysate between the HepG2 cells treated with different EVs or rHuArg1 at the same time points. A p value <0.05 was considered statistically significant. For in vivo studies, a 2-tailed t-test or a one-way ANOVA controlled for multiple comparisons were used. Details are to be found in figure legends. All the statistical analyses were performed using GraphPad Prism Version 10.

## Authors Contribution

LEH, MC, MS designed the studies. MS designed the constructs. LEH performed the in vitro studies. MC, BM performed the in vivo studies. CL, MJL, WM produced the exosomes. LEH, MC, MS wrote the manuscript.

## Disclosure

All authors are employees at Capricor Therapeutics.

## Supplementary Figures

**Supplementary Figure 1.**
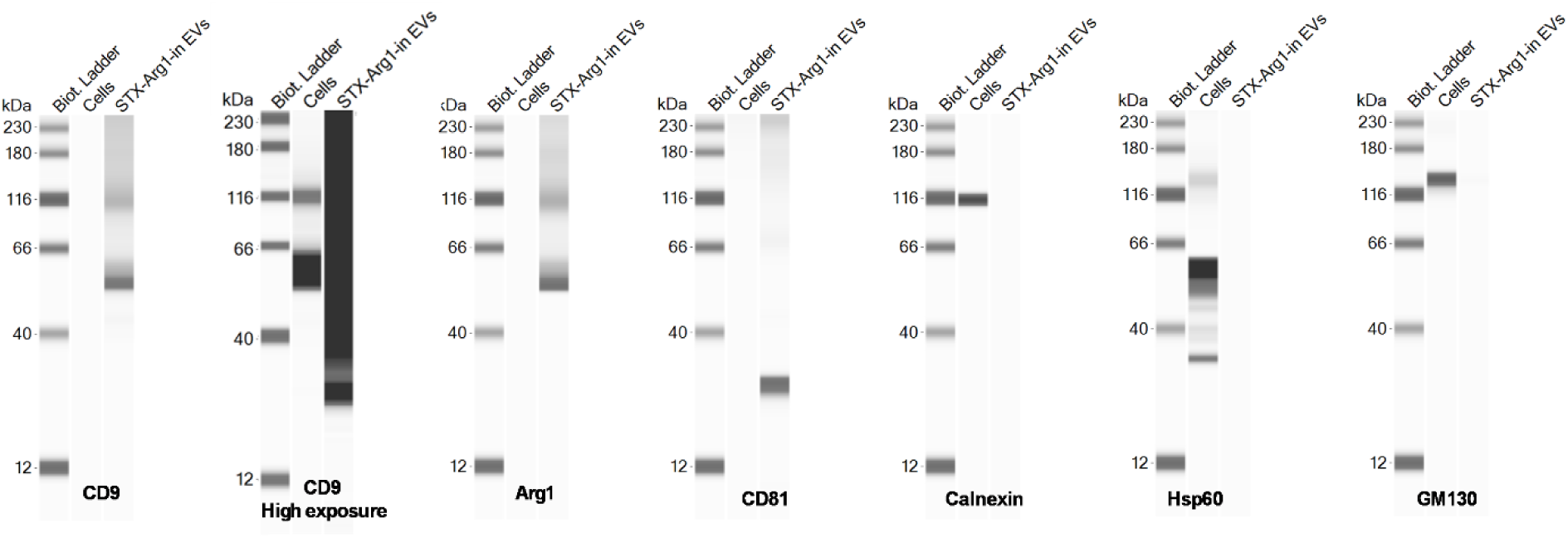
Jess western blot. Representative image of Cells and STX-Arg1-in EVs probed for exosome markers (CD9, CD81), Arginase 1 (ARG1), cell contamination markers (calnexin, Hsp60) and Golgi marker (GM130).

**Supplementary Figure 2.**
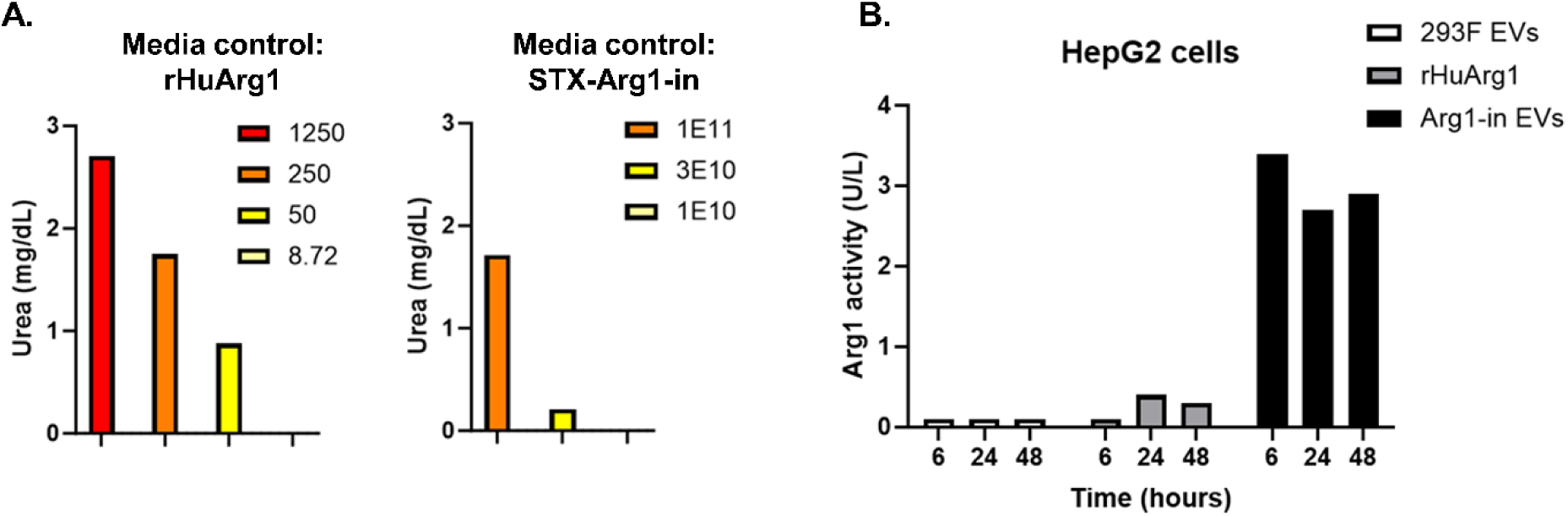
**A.** Activity of rHuArg1 and STX-Arg1-in EVs in absence of cells**. B.** 293F EVs have no biological effects in HepG2 cells. HepG2 cells were treated with 293F EVs, recombinant human Arg1 (rHuArg1) and Arg1-in EVs and ARG1 activity measured in cells at 6, 24 and 48 hours after treatment. No activity was observed with 293F EVs treatment, or rHuARG1, compared to STX-ARG1-in EVs. N=3.

**Supplementary Figure 3.**
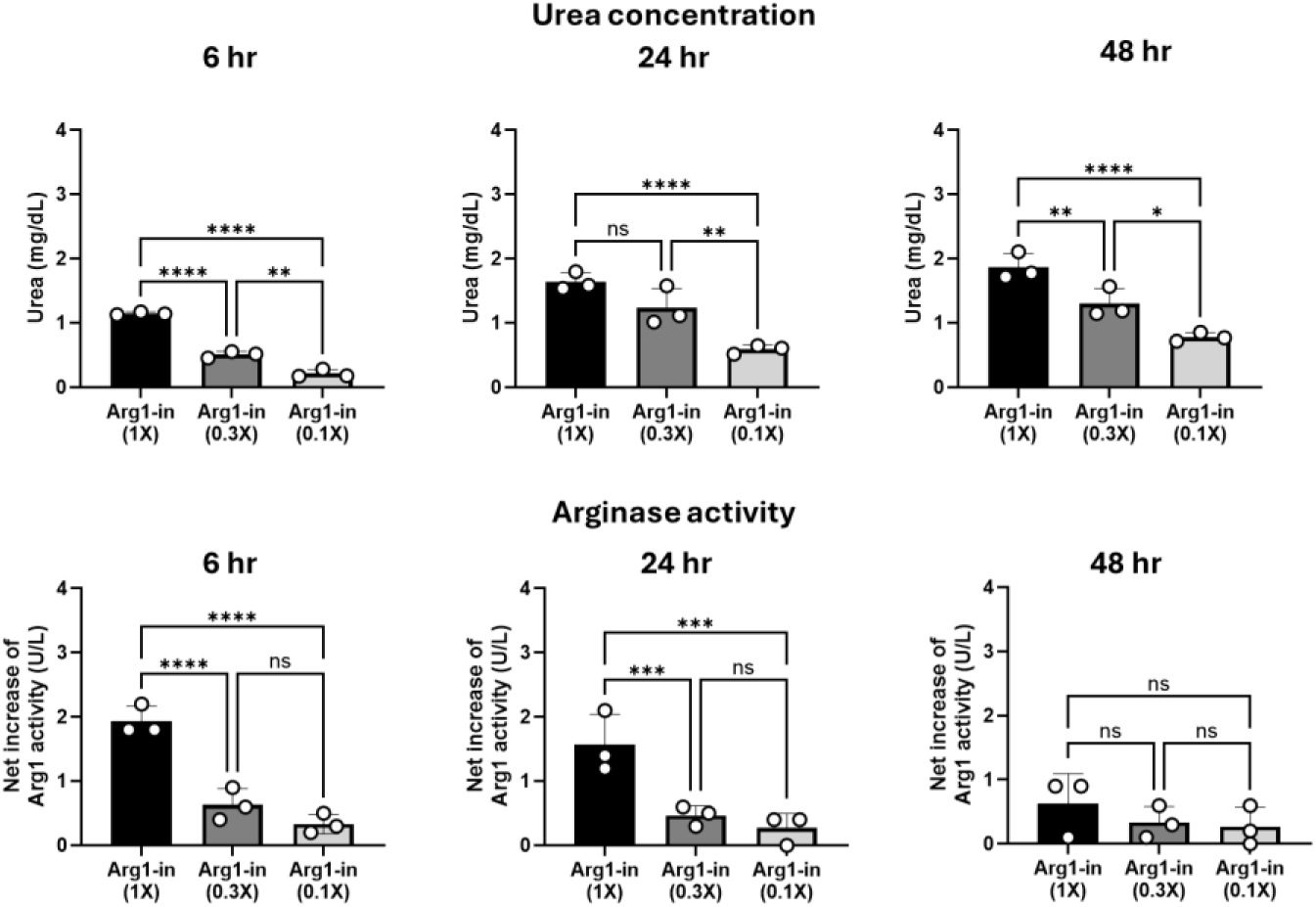
Dose response of STX-Arg1-EVs: urea production and ARG1 activity. HepG2 cells were treated with 1X (1E11 EVs), 0.3X (3E10 EVs) and 0.1X (1E10 EVs) dose of STX-Arg1-in EVs to identify the optimal dose (EVs amount, ARG1 concentration). After treatment, urea concentration and arginase activity were analyzed and quantified at 6, 24 and 48 hours. Data was analyzed using an ordinary one-way ANOVA with Tukey’s multiple comparison test. A p value < 0.05 was considered to be statistically significant. ns: not significant. *: P < 0.05. **: P < 0.01. ***: P < 0.001. ****: P < 0.0001.

**Supplementary Figure 4.**
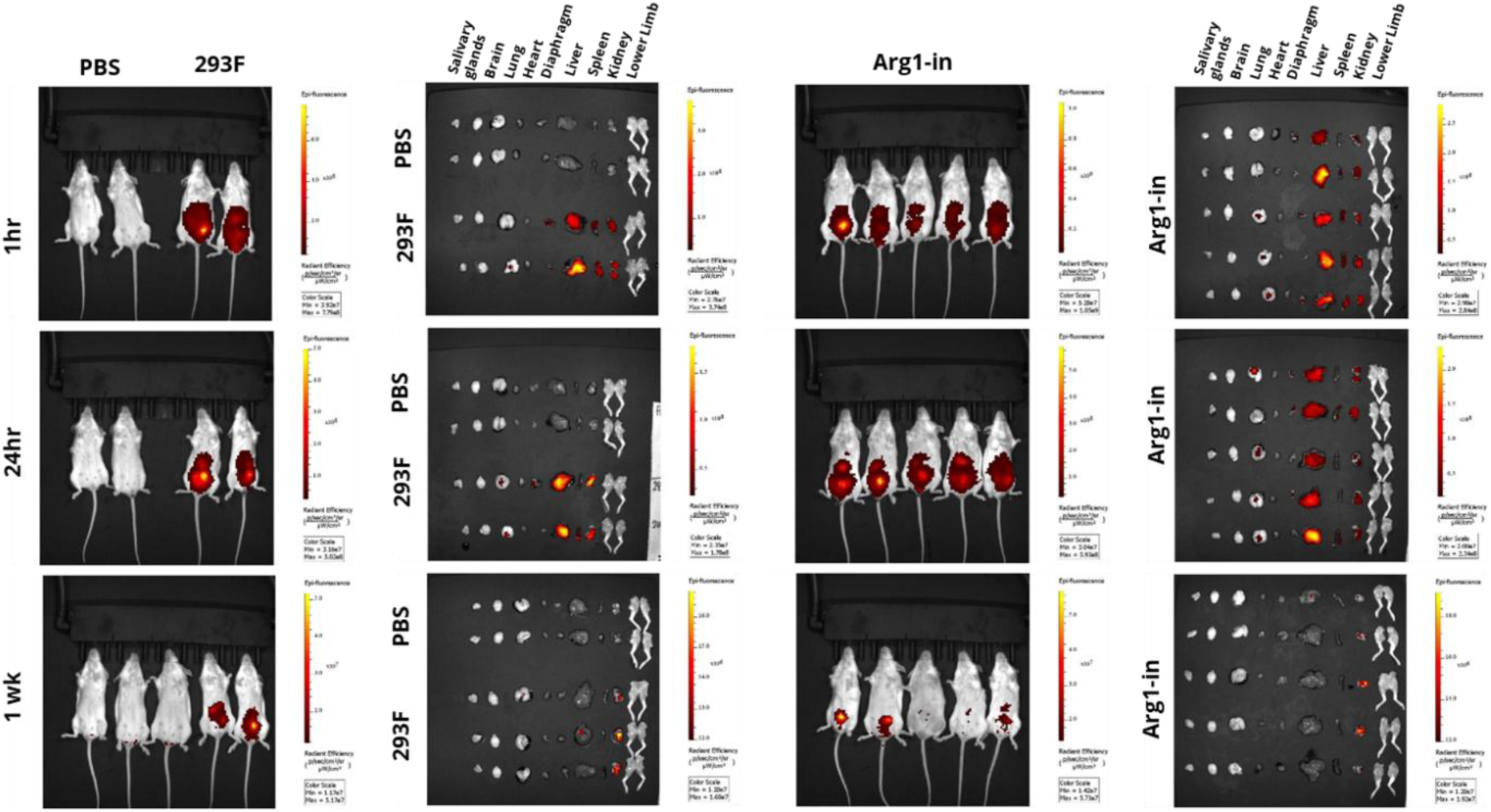
In vivo imaging by IVIS imager for exosome biodistribution. Representative images of full body and isolated tissues from wild-type mice receiving intraperitoneal injection of labeled STX-Arg1-EVs at 1hr, 24hr and 1 week (wk) after injection..

**Supplementary Figure 5.**
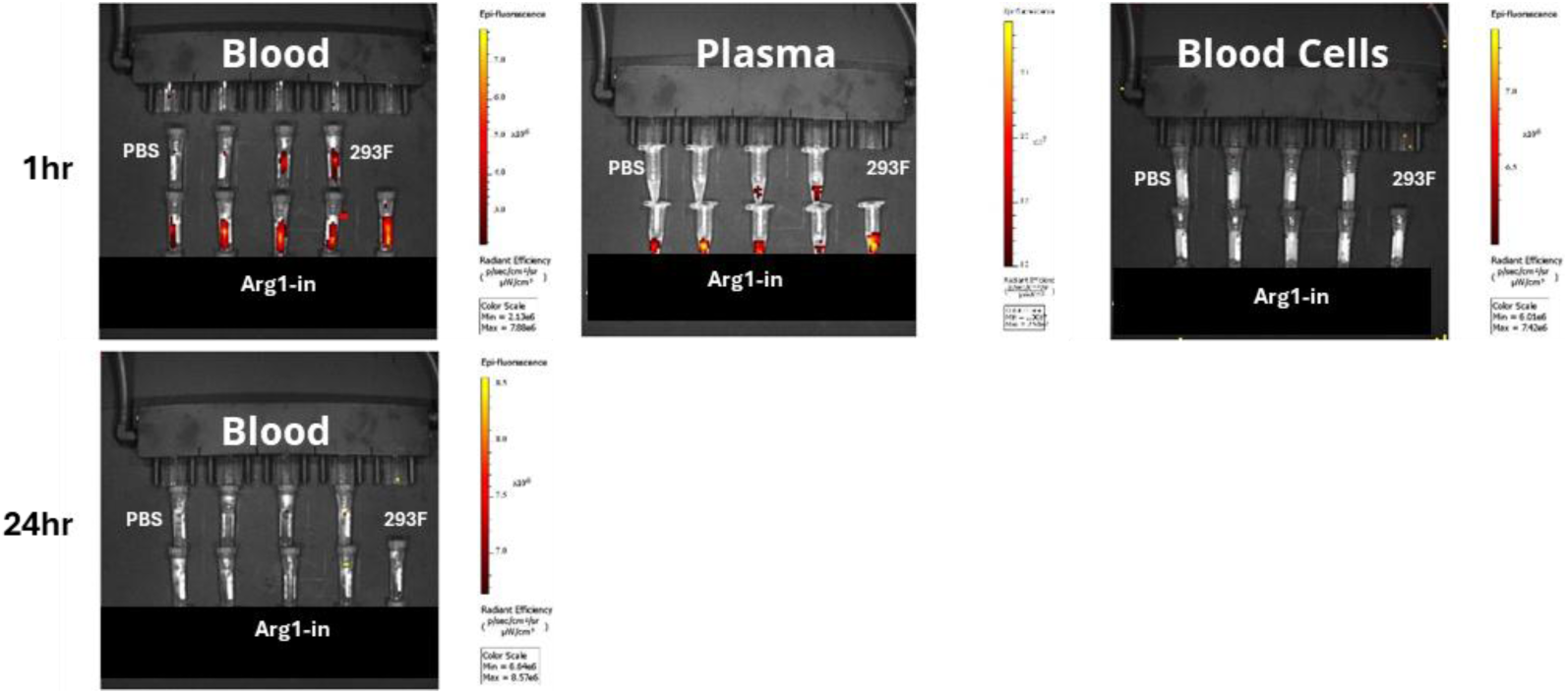
IVIS imaging of blood. Representative images of full blood, plasma and blood cells from wild-type mice receiving intraperitoneal injection of labeled STX-Arg1-EVs at 1hr and 24hr after injection..

**Supplementary Figure 6.**
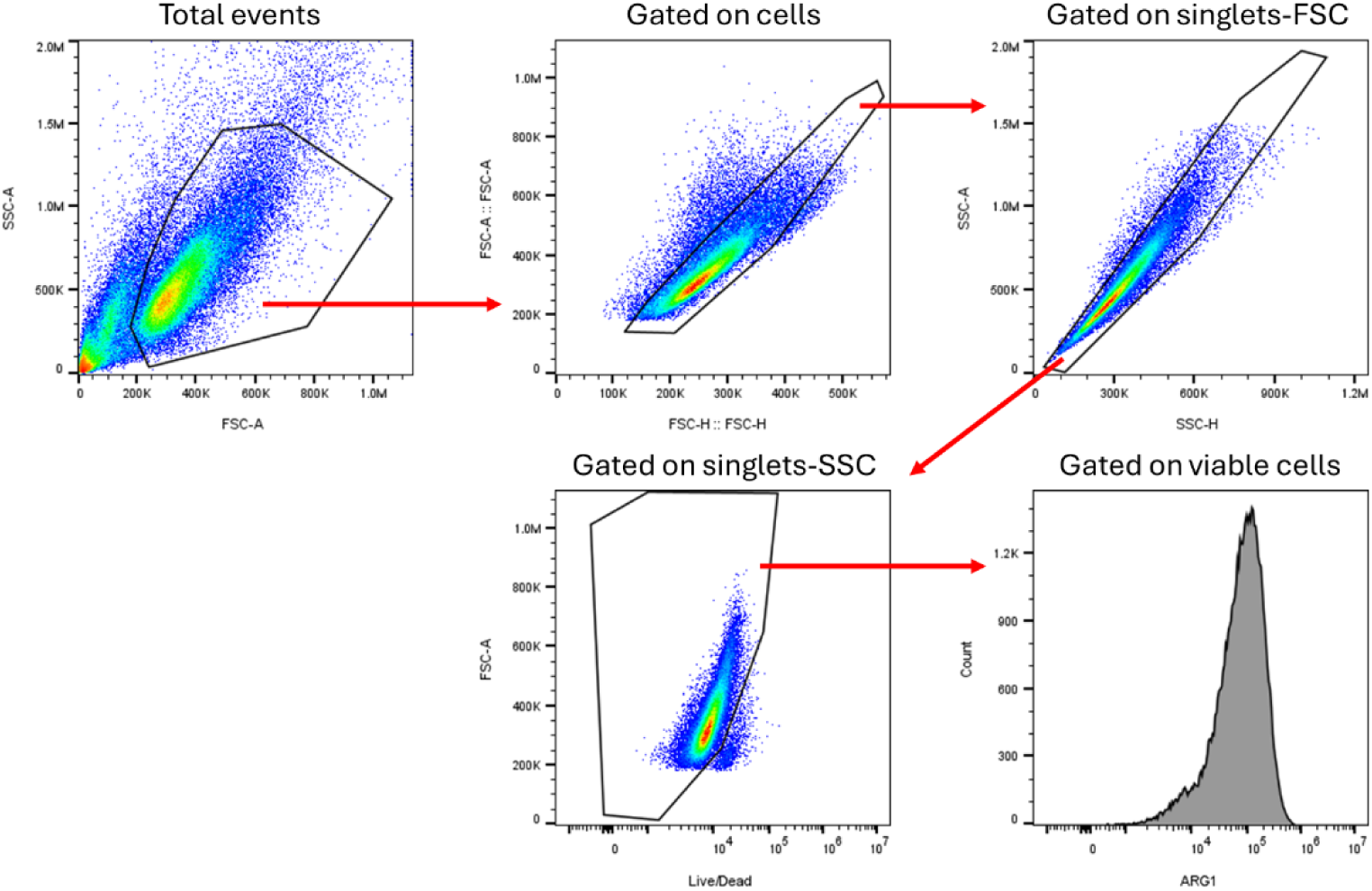
Gating strategy to measure ARG1 expression of engineered cell lines. Representative image of gating strategy for quantification of ARG1 expression of engineered cell lines.

